# Cardenolides May Affect Herbivory on Milkweeds (*Asclepias* spp.) by the Root-Knot Nematode *Meloidogyne incognita*

**DOI:** 10.1101/2025.06.30.662457

**Authors:** Damaris Godinez-Vidal, Perla Achi, Adler R. Dillman, Simon C. Groen

## Abstract

Root-knot nematodes (RKNs) of the genus *Meloidogyne* are important pests in agriculture. RKNs are generalist herbivores with a wide host range including crop and wild plants. The latter are an important source of defensive metabolites that may be helpful for RKN management. Milkweeds (*Asclepias* spp.) produce toxic cardenolides that protect them from herbivory. However, it is unclear if cardenolides may defend milkweeds against RKNs. Here, we tested this directly through herbivory assays with the RKN *M. incognita* on milkweed species that produce negligible and high levels of cardenolides, *A. tuberosa* and *A. curassavica*, respectively. We found that *M. incognita* induces fewer galls on *A. curassavica* than *A. tuberosa* and fails to reproduce after reaching maturity on the former but not the latter species. This suggests that the predominantly polar cardenolides in *A. curassavica* may engender long-term reproductive toxicity. Further toxicity assays with the polar cardenolide ouabain showed that cardenolides can also have more immediate toxic effects on *M. incognita* at higher concentrations. Ouabain caused a coiled, paralytic phenotype in juvenile RKNs, a sign of neurotoxicity, leading to lethality in a subset of RKNs. Some nematodes recovered upon ouabain removal, confirming that neurotoxic cardenolides have a nematostatic effect that results in death when exposure persists. Taken together, our results provide further evidence that cardenolides may function as anti-herbivore defenses against RKNs. The ‘dead end’ host plant *A. curassaviva* appears to possess useful properties that may be leveraged for control of RKNs in agriculture.

## Introduction

RKNs are plant-parasitic nematodes that attack a wide variety of plants and cause significant yield losses in numerous agricultural crops, estimated at 10–20% yield reduction (Koenning et al., 1999). Crop losses from thermophilic RKNs such as *M. incognita* can be particularly severe in tropical and subtropical regions where their reproductive rates are generally higher than in temperate regions. Yield reductions caused by RKNs might exacerbate in the future due to restrictions imposed on the use of certain classes of nematicides (Hooks et al., 2010). There is therefore much interest in the development of alternative approaches for managing RKNs, including adoption of nematode-resistant or -antagonistic plants in agricultural systems. One example is incorporation of marigold (*Tagetes* spp.), which is resistant to RKNs through the production of the anti-herbivore defensive metabolite alpha-terthienyl (Hooks et al., 2010). Another is the use of litchi tomato (*Solanum sisymbriifolium*), which may derive its toxicity to RKNs from the production of glycoalkaloids such as α-solamargine (Baker et al., 2023; Pillai and Dandurand, 2021; Udalova et al., 2004). A particularly useful feature of these plants is that they may serve as ‘dead end’ hosts or ‘trap crops’: eggs may hatch in the presence of these plants and juvenile RKNs may even infect them, but then the nematodes ultimately perish due to the plants’ toxicity. Consequently, significant reductions of RKN population densities might occur when such plants are incorporated in RKN management practices (Baker et al., 2023; Hooks et al., 2010).

Milkweeds in the Pan-American genus *Asclepias* form another potential source of RKN control. These plants produce toxic anti-herbivore defensive metabolites known as cardenolides, which are cardiac glycosides with a five-membered lactone group (Agrawal et al., 2012). Cardiac glycosides inhibit the essential animal enzyme Na^+^/K^+^-ATPase and can have potent neurotoxic effects on herbivores that have not adapted to feeding on milkweeds (Groen et al., 2017; Groen and Whiteman, 2022). Potential toxic effects of milkweeds on RKNs have been studied for several combinations of species. The RKN *M. incognita* has been found to induce gall formation when infecting *A. curassavica* (Haseeb and Pandey, 1987; Lima-Medina et al., 2013; López and Quesada, 1997). However, the nematode’s reproduction factor was close to zero (0.014) and gall formation was no longer observed when juveniles from the subsequent generation were inoculated onto a susceptible tomato host, suggesting a toxic effect of *A. curassavica* that could become apparent at later stages of *M. incognita*’s life cycle (López and Quesada, 1997). Similar toxic effects were observed for the RKN *M. chitwoodi* when soils were amended with seedmeal of *A. speciosa* and *A. syriaca* as this led to an inability of nematodes to infect roots of bioassay tomato plants. However, additional experiments revealed that the decline of *M. chitwoodi* in amended soil may not have been due to cardenolides but more likely to the presence of other soluble natural products in milkweed seedmeal (Harry-O’Kuru et al., 1999).

Further experiments with *M. incognita* infections of *A. fascicularis* and *A. speciosa* showed that this nematode induced almost twice as many galls on *A. speciosa* plants than on *A. fascicularis* plants, despite both species showing similar cardenolide levels (Mundim and Pringle, 2020). However, *A. fascicularis* plants exhibited rotting and abortion of infected root parts. Furthermore, *M. incognita* infection reduced both shoot and root growth of *A. fascicularis* and ultimately reduced plant survival in this species. Although *M. incognita* infection did not affect the survival of *A. speciosa* plants, it did reduce relative shoot growth in this species (Mundim and Pringle, 2020). Taken together, it is unclear from these studies if cardenolides could play a role in limiting *M. incognita* infection of milkweeds.

To test if cardenolides might affect milkweed infection by *M. incognita*, we compared infections by this nematode on *A. curassavica*, a milkweed species that produces relatively high levels of cardenolides, and *A. tuberosa*, a milkweed that almost completely lacks cardenolides (Rasmann and Agrawal, 2011a). Cardenolides from *A. curassavica* tend to have a relatively high polarity index (Rasmann and Agrawal, 2011b), and we therefore performed *in vitro* toxicity tests with the polar cardenolide ouabain to confirm if these compounds could affect *M. incognita*.

## Materials and Methods

### Infection of milkweeds with *Meloidogyne incognita*

Seeds of the milkweed species *Asclepias curassavica* and *A. tuberosa* (W. Atlee Burpee and Co., Warminster, PA, USA) were germinated in peat moss for six days until the first true leaf was observed. Seedlings were transplanted in soil mix (90% sand, 10% peat moss) and maintained in pots (5 cm × 6 cm × 8.5 cm) in a plant growth room with a 16-hour photoperiod at 28 ºC. For the RKN infection experiment, eggs from *M. incognita* population Project 77 Race 3 (P77R3 / VW6) were extracted from infected roots of tomato (*Solanum lycopersicum*) cultivar Moneymaker (Godinez-Vidal et al., 2024). Eggs were floated on a 35% sucrose cushion and cleaned with 10% bleach. Eggs were hatched in a glass dish with a mesh insert and tissue paper at 28°C in the dark for five days. The number of second-stage juvenile (J2) nematodes that hatched was quantified and a nematode inoculum suspension was prepared. Two-week-old milkweed seedlings, with three true leaves, were inoculated with 200 and 500 J2-stage nematodes. Root tissue was collected at 1, 3, 6, 12, 17, 21, 27, and 39 days after inoculation (dai). The roots were rinsed and cleared with 10% bleach for 5 min, then stained with 1× acid fuchsin (3.5 g acid fuchsin, 125 mL acetic acid, 375 mL H_2_O) and preserved in 50% glycerol at room temperature. The roots were placed on coverslips and observed with an optical microscope. Images of roots were obtained using a Leica Microscope, objective 10×/0.5 Plan M, with a Nikon DS Camera Head, and Nikon NIS Elements Imaging Software to process images. The experiment was performed in triplicate.

### In vitro cardenolide toxicity assay on *Meloidogyne incognita*

The effects of the purified cardenolide ouabain were evaluated *in vitro* on J2-stage nematodes of *M. incognita* at different exposure times (Godinez-Vidal and Groen, 2025). Ouabain (Millipore Sigma) was dissolved in distilled water, and a stock solution was prepared. After hatching, 100 J2-stage RKNs were set up in a microsyracuse glass container with solutions of different ouabain concentrations: 0 mM (distilled water control), 5 mM, and 15 mM. The effects of ouabain on nematodes were evaluated at 24, 48, and 72 hours of exposure. After 72 hours, the ouabain was removed, and distilled water was added to the microsyracuse glass containers. The experiment was performed at 28 ºC in dark conditions. Phenotyping observations were performed at 24 hours after adding distilled water and the numbers of coiled-up or dead nematodes were quantified. Images of roots were obtained using a Leica Microscope, objective 10×/0.5 Plan M, with a Nikon DS Camera Head. The experiment was performed in triplicate.

*Graphing and statistical analysis*. All graphs were made using PRISM with the mean and standard error of the mean (SEM). Statistical analysis was performed using one-way or two-way ANOVA followed by post-hoc Tukey’s HSD tests depending on the data analyzed. Figures were assembled using Adobe Photoshop.

## Results

### *Meloidogyne incognita* fails to reproduce on high-cardenolide *Asclepias curassavica* plants

To evaluate and compare infections of *A. curassavica* and *A. tuberosa* roots by *M. incognita*, we inoculated seedlings of both *Asclepias* species with 200 and 500 J2-stage nematodes and collected the inoculated roots at 1, 3, 6, 12, 17, 21, 27, and 39 dai. The presence of J2-stage nematodes in root tips of the plants was observed at 3 dai, with the highest nematode numbers observed at 6 dai in both species (P<0.001), whether they had been inoculated with 200 (Fig. 1a and b) or 500 nematodes (Fig. 1c and d). At 17 dai, the presence of nematodes in the next stages of the life cycle, J3-J4, became apparent and their numbers were similar in both *Asclepias* species. However, at 27 dai the number of females in roots was significantly higher in *A. tuberosa* than in *A. curassavica* (P<0.001; Fig. 1d), suggesting that *M. incognita* had better access to host resources in *A. tuberosa*. This finding was corroborated by the observation of significantly higher numbers of galls in *A. tuberosa* compared to *A. curassavica* at 17 dai (P<0.001), a difference that was maintained up to 39 dai (Fig. 1e and f). We further observed that females of *M. incognita* were able to produce egg masses in *A. tuberosa* but not in *A. curassavica* (Fig. 2). This suggests that in the long term, *M. incognita* is less affected by the defenses of *A. tuberosa*, which does not produce cardenolides, than it is by the defenses of *A. curassavica*, which produces relatively high levels of cardenolides, affecting the nematodes’ ability to reproduce.

**Fig. 1.**
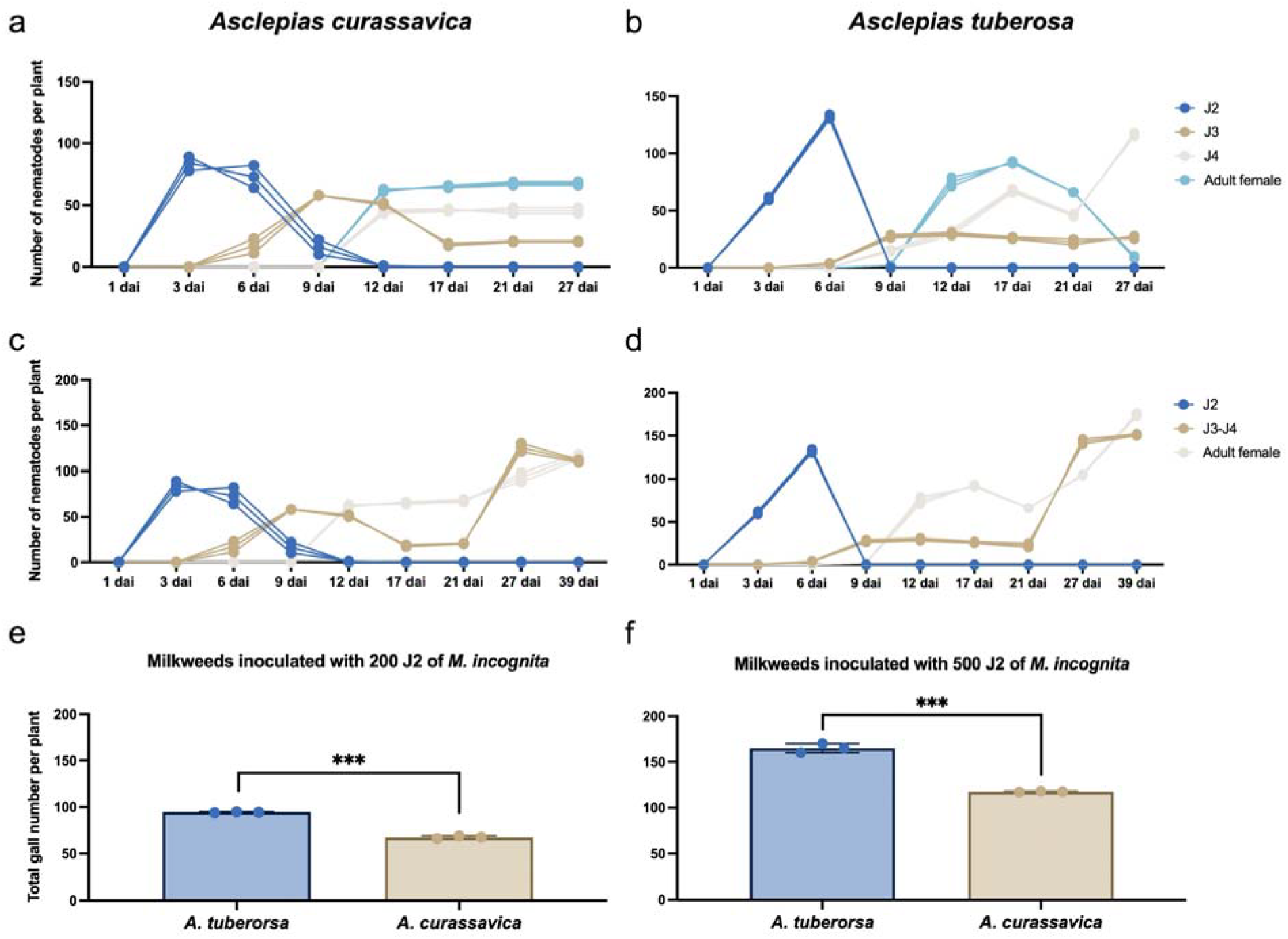
*Meloidogyne incognita* can infect roots of the butterfly milkweed *Asclepias tuberosa* and the tropical milkweed *A. curassavica*. a. Number of *M. incognita* nematodes at different life stages over time in *A. curassavica* inoculated with 200 J2-stage nematodes of *M. incognita* population P77R3 / VW6. b. Results from the same experiments as in panel a for *A. tuberosa*. c. Number of *M. incognita* nematodes at different life stages over time in *A. curassavica* inoculated with 500 J2-stage nematodes. d. Results from the same experiments as in panel a for *A. tuberosa*. e. Number of galls in *Asclepias* spp. plants at 39 days after inoculation with 200 J2-stage nematodes. f. Results from the same time point as in panel e after inoculation with 500 J2-stage nematodes. Asterisks in e and f indicate P values <0.001 (two-way ANOVA with post-hoc Tukey’s HSD tests).

**Fig. 2.**
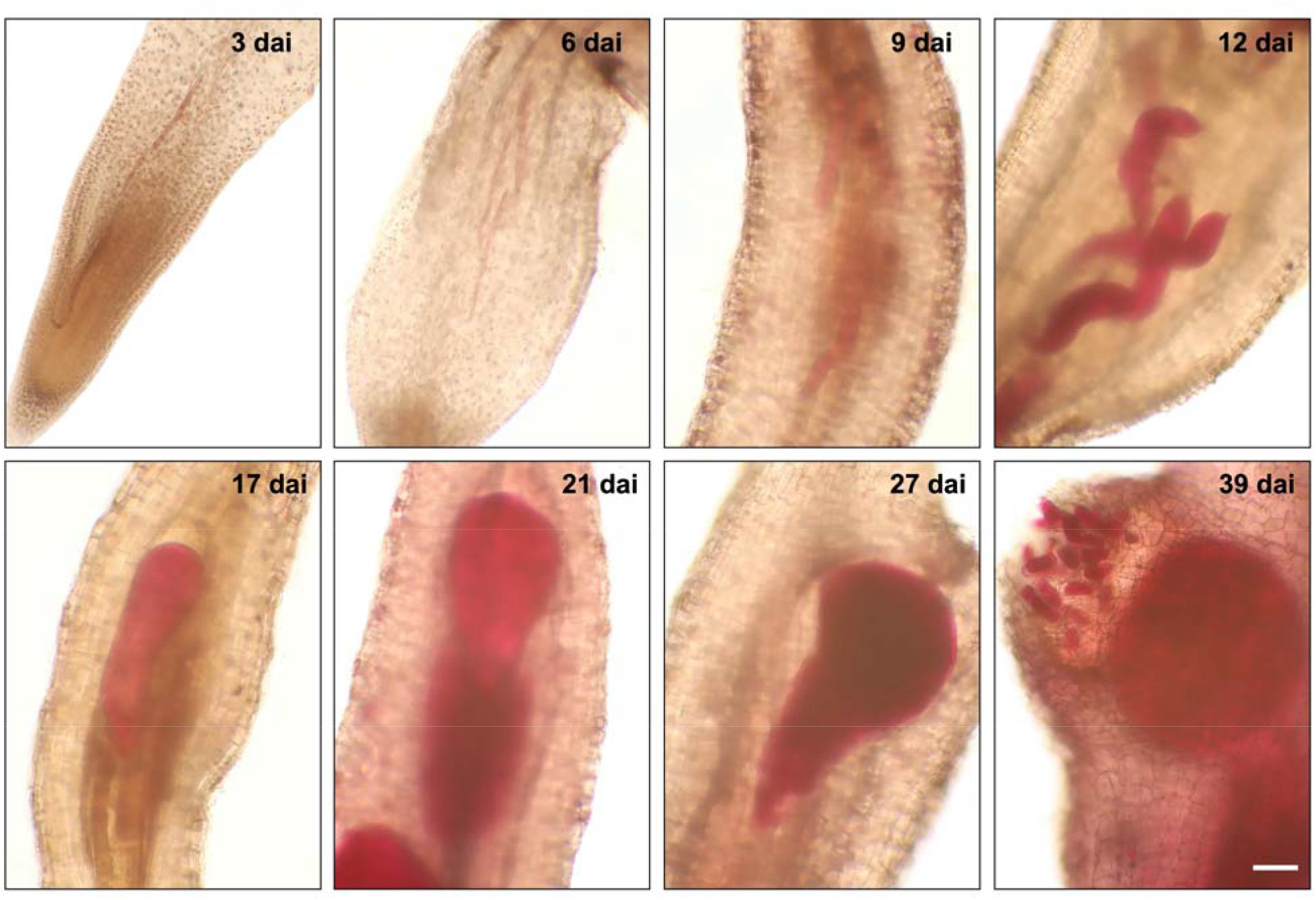
*Meloidogyne incognita* can reproduce in roots of *Asclepias tuberosa* but not in roots of *A. curassavica*. Representative images of different life stages of *M. incognita* population P77R3 / VW6 were selected for nematodes in roots of *A. tuberosa* at 3, 6, 9, 12, 17, 21, 27, and 39 days after inoculation (dai). The image taken at 39 dai shows an egg mass produced by an adult female nematode (scale bar = 200 μm).

### Cardenolides can have neurotoxic effects on *Meloidogyne incognita* juveniles

To confirm the sensitivity of *M. incognita* to cardenolides, we evaluated the shorter-term effects of higher levels of the purified polar cardenolide ouabain on J2-stage RKNs at different exposure times in an *in vitro* toxicity assay. At 24 hours of exposure, we observed a coiled phenotype (curved body posture in one direction) when RKNs were exposed to the relatively high ouabain concentration of 15 mM (Figure 3a). This coiled phenotype is a known sign of a compound’s neurotoxic effects on nematodes (Harrington et al. 2023). However, the coiled phenotype was also observed after exposure to the lower concentration of 5 mM at 48 and 72 hours after the start of the assay (Figure 3a). This suggests that lower concentrations of ouabain can affect *M. incognita* but that exposure time is important for juvenile nematodes to become sensitive to the toxin. We confirmed this observation at 72 hours of exposure, where both the lower and higher concentrations of ouabain had a toxic effect on J2-stage RKNs (Figure 3b), indicating that the effects of different ouabain concentrations are directly proportional to the time of exposure. At 72 hours of exposure, we further observed RKN death (straight body posture) with both concentrations of ouabain (Figure 3c and d). To confirm the neurotoxic effect of ouabain, we removed it at 72 hours of exposure and switched the J2-stage RKNs to distilled water for 24 hours. We observed that some RKNs maintained the coiled phenotype or reverted to a normal body posture, which showed that ouabain had a nematostatic effect. However, 40% and 50% of RKNs showed lethality when exposed to 5 mM and 15 mM ouabain, respectively (Figure 3c and d), suggesting that ouabain’s nematostatic effect engendered a nematicidal effect on *M. incognita* juveniles after prolonged exposure.

**Fig. 3.**
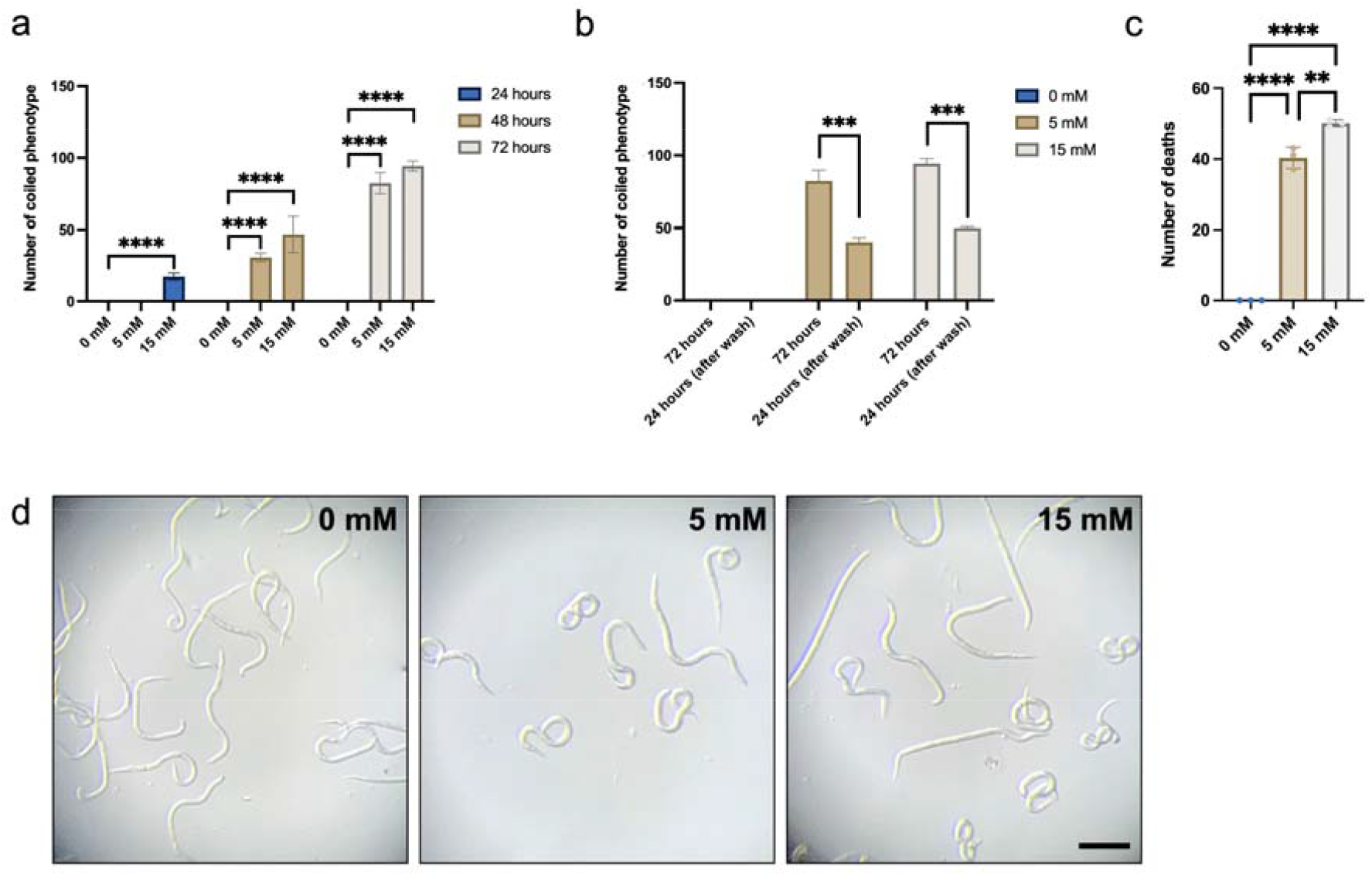
The polar cardenolide ouabain has neurotoxic effects on *Meloidogyne incognita*. a. Number of J2-stage *M. incognita* nematodes exhibiting a coiled, paralytic phenotype at 24, 48, and 72 hours of exposure to 0 mM, 5 mM or 15 mM of ouabain. b. Number of J2-stage *M. incognita* nematodes exhibiting a coiled, paralytic phenotype at 72 hours of exposure to 0 mM, 5 mM or 15 mM of ouabain before and after a subsequent washing step with distilled water for 24 hours. c. Number of J2-stage *M. incognita* nematodes exhibiting a lethal phenotype at 72 hours of exposure to 0 mM, 5 mM or 15 mM of ouabain. d. J2-stage *M. incognita* nematodes displaying a coiled phenotype (curved body posture in one direction) and death phenotype (straight body posture) at 72 hours of exposure to 0 mM, 5 mM or 15 mM of ouabain (scale bar = 50 μm).

## Discussion

In this study we found evidence that cardenolides may have long- and short-term toxic effects on *M. incognita* RKNs. Comparing nematode life cycles on milkweed plants with relatively high levels of cardenolides (*A. curassavica*) versus life cycles on milkweed plants almost entirely devoid of cardenolides (*A. tuberosa*) revealed that *M. incognita* was able to complete its life cycle on *A. tuberosa* plants but not on cardenolide-rich *A. curassavica* plants. This difference in toxicity was further evident from the observation that J2-stage RKNs induced more galls on *A. tuberosa* than on *A. curassavica*.

The results for *A. curassavica* reveal useful features of this plant as a potential trap crop for use in RKN management. Trap crops such as litchi tomato and marigolds may promote nematode egg hatching and allow juveniles to even infect the plants, but then intoxicate the nematodes by producing potent defensive chemicals such as glycoalkaloids and alpha-terthienyl (Baker et al., 2023; Hooks et al., 2010). The tropical milkweed *A. curassavica* appears to have similar effects on RKNs: J2-stage *M. incognita* nematodes readily infected this species—albeit to a somewhat lesser extent than the low-cardenolide milkweed *A. tuberosa*—but while the nematodes were able to continue development on *A. curassavica* until the adult female stage, these females were unable to produce egg masses on this high-cardenolide milkweed. These results align with previous observations of *M. incognita*’s reproductive fitness on *A. curassavica* in greenhouse conditions (López and Quesada, 1997). However, more greenhouse and field studies will be necessary to further explore the use of this plant in soil amendments, as a cover crop or in intercropping practices (Harry-O’Kuru et al., 1999; Hooks et al., 2010).

In addition to potential long-term toxic effects of milkweed-produced cardenolides on *M. incognita*, our *in vitro* toxicity assays revealed that higher levels of cardenolides can further have more immediate neurotoxic effects on this nematode. Exposure to millimolar levels of the polar cardenolide ouabain led to a coiled, paralytic phenotype in J2-stage RKNs, which is a sign of nematostatic effects and neurotoxicity (Harrington et al., 2023). Longer exposure ultimately had a nematicidal effect and led to lethality among the nematodes. Similar toxic effects have previously been observed for entomopathogenic nematodes of the genus *Steinernema* and the free-living nematode *Caenorhabditis elegans* (Achi et al., 2025; Davis et al., 1995). A large body of research has shown that cardenolides exert their neurotoxicity by inhibiting the Na^+^/K^+^-ATPase, which is ubiquitously expressed in animals (Agrawal et al., 2012). This includes expression in their nervous systems and cardenolides prevent nematodes from engaging in normal feeding behavior by blocking the Na^+^/K^+^-ATPase (Davis et al., 1995).

One way through which several milkweed-specialized herbivores such as the monarch butterfly have adapted to the presence of toxic cardenolides in their diets is evolution of amino acid substitutions in the first extracellular loop of the Na^+^/K^+^-ATPase that confer target-site insensitivity and resistance to cardenolides (Karageorgi et al., 2019). Nematodes, including RKNs, share amino acid substitutions at two sites (111 and 119) in the first extracellular loop with some of these specialist herbivores and it will be interesting to study if the amino acid residues RKNs carry at these sites, D111 and S119, confer a somewhat lowered sensitivity to cardenolides in RKNs relative to insect herbivores (Groen and Whiteman, 2021).

In summary, our experiments uncovered evidence for toxic effects of cardenolides, and of milkweeds that produce these defensive metabolites, on *M. incognita*. These findings lay the foundation for future explorations of how milkweeds may be leveraged for management of RKNs in agricultural crop production systems.

## Acknowledgements

The authors wish to thank the National Institute of General Medical Sciences of the National Institutes of Health (award no. R35GM151194 to SCG and award no. R35GM137934 to ARD), the United States Department of Agriculture’s National Institute of Food and Agriculture (award no. 1032189 to ARD), and the University of California Riverside (start-up funds to SCG) for support.

## References

Achi, P., Christensen, P., Iglesias, V., McCarthy, C., Pena, R., Bavier, L., Goldy, C., Agrawal, A.A., Groen, S.C., and Dillman, A.R. 2025. Entomopathogenic nematode species vary in their behavior and virulence in response to cardiac glycosides within and around insect hosts. Journal of Chemical Ecology 51:12.

Agrawal, A.A., Petschenka, G., Bingham, R.A., Weber, M.G., and Rasmann, S. 2012. Toxic cardenolides: chemical ecology and coevolution of specialized plant-herbivore interactions. New Phytologist 194:28–45.

Baker, H.V., Zasada, I.A., Gleason, C., and Dandurand, L.M. 2023. Reproduction and invasion dynamics of Globodera pallida and Meloidogyne spp. in Solanum sisymbriifolium. Nematropica 53:70–81.

Davis, M.W., Somerville, D., Lee, R.Y., Lockery, S., Avery, L., and Fambrough, D.M. 1995. Mutations in the Caenorhabditis elegans Na, K-ATPase alpha-subunit gene, eat-6, disrupt excitable cell function. Journal of Neuroscience 15:8408–8418.

Godinez-Vidal, D., Edwards, S.M., and Groen, S.C. 2024. Root-knot nematode egg extraction. protocols.io 10.17504/protocols.io.eq2lyj5nqlx9/v1.

Godinez-Vidal, D., and Groen, S.C. 2025. In vitro compound toxicity protocol for nematodes. protocols.io 10.17504/protocols.io.n2bvj9bxplk5/v1.

Groen, S. C., LaPlante, E. R., Alexandre, N. M., Agrawal, A. A., Dobler, S., and Whiteman, N. K. 2017. Multidrug transporters and organic anion transporting polypeptides protect insects against the toxic effects of cardenolides. Insect Biochemistry and Molecular Biology 81:51–61.

Groen, S.C., and Whiteman, N.K. 2021. Convergent evolution of cardiac-glycoside resistance in predators and parasites of milkweed herbivores. Current Biology 31:R1465–R1466.

Groen, S.C., and Whiteman, N.K. 2022. Ecology and evolution of secondary compound detoxification systems in caterpillars. In: Caterpillars in the Middle: Tritrophic Interactions in a Changing World (pp.115–163). Springer International Publishing.

Harrington, S., Pyche, J., Burns, A.R., Spalholz, T., Ryan, K.T., Baker, R.J., Ching, J., Rufener, L., Lautens, M., Kulke, D., Vernudachi, A., Zamanian, M., Deuther-Conrad, W., Brust, P., and Roy, P.J. 2023. Nemacol is a small molecule inhibitor of C. elegans vesicular acetylcholine transporter with anthelmintic potential. Nature Communications 14:1816.

Harry-O’Kuru, R.E., Mojtahedi, H., Vaughn, S.F., Dowd, P.F., Santo, G.S., Holser, R.A., and Abbott, T.P. 1999. Milkweed seedmeal: a control for Meloidogyne chitwoodi on potatoes. Industrial Crops and Products 9:145–150.

Haseeb, A., and Pandey, R. 1987. Incidence of root-knot nematodes in medicinal and aromatic plants – new host records. Nematropica 17:209–212.

Hooks, C.R., Wang, K.H., Ploeg, A.T., and McSorley, R. 2010. Using marigold (Tagetes spp.) as a cover crop to protect crops from plant-parasitic nematodes. Applied Soil Ecology 46:307–320.

Karageorgi, M., Groen, S.C., Sumbul, F., Pelaez, J.N., Verster, K.I., Aguilar, J.M., Hastings, A.P., Bernstein, S.L., Matsunaga, T., Astourian, M. and Guerra, G., Rico, F., Dobler, S., Agrawal, A.A., and Whiteman, N.K. 2019. Genome editing retraces the evolution of toxin resistance in the monarch butterfly. Nature 574:409–412.

Lima-Medina, I., Somavilla, L., Carneiro, R.M.D.G., and Gomes, C.B. 2013. Species of Meloidogyne associated with fig (Ficus carica) and host weeds. Nematropica 43:56–62.

López, R., and Quesada, M. 1997. Reproduction of Meloidogyne incognita on several weeds present in Costa Rica. Agronomía Mesoamericana 8:112–115.

Mundim, F.M., and Pringle, E.G. 2020. Phytochemistry-mediated disruption of ant–aphid interactions by root-feeding nematodes. Oecologia 194:441–454.

Pillai, S.S., and Dandurand, L.M. 2021. Effect of steroidal glycoalkaloids on hatch and reproduction of the potato cyst nematode Globodera pallida. Plant Disease 105:2975–2980.

Rasmann, S., and Agrawal, A.A. 2011a. Evolution of specialization: a phylogenetic study of host range in the red milkweed beetle (Tetraopes tetraophthalmus). The American Naturalist 177:728–737.

Rasmann, S., and Agrawal, A.A. 2011b. Latitudinal patterns in plant defense: evolution of cardenolides, their toxicity and induction following herbivory. Ecology Letters 14:476–483.

Udalova, Z.V., Zinov’eva, S.V., Vasil’eva, I.S., and Paseshnichenko, V.A. 2004. Correlation between the structure of plant steroids and their effects on phytoparasitic nematodes. Applied Biochemistry and Microbiology 40:93–97.

